# An acute immune response underlies the benefit of cardiac adult stem cell therapy

**DOI:** 10.1101/506626

**Authors:** Ronald J. Vagnozzi, Marjorie Maillet, Michelle A. Sargent, Hadi Khalil, Anne Katrine Johansen, Jennifer A. Schwanekamp, Allen J. York, Vincent Huang, Matthias Nahrendorf, Sakthivel Sadayappan, Jeffery D. Molkentin

## Abstract

Clinical trials using adult stem cells to regenerate damaged heart tissue continue to this day^1–3^ despite ongoing questions of efficacy and a lack of mechanistic understanding of the underlying biologic effect^4–6^. The rationale for these cell therapy trials is derived from animal studies that show a modest but reproducible improvement in cardiac function in models of cardiac ischemic injury^7–9^. Here we examined the mechanistic basis for cell therapy in mice after ischemia/reperfusion (I/R) injury, and while heart function was enhanced, it was not associated with new cardiomyocyte production. Cell therapy improved heart function through an acute sterile immune response characterized by the temporal and regional induction of CCR2^+^ and CX3CR1^+^ macrophages. Here we observed that intra-cardiac injection of 2 distinct types of progenitor cells, freeze/thaw-killed cells or a chemical inducer of the innate immune response similarly induced regional CCR2^+^ and CX3CR1^+^ macrophage accumulation and provided functional rejuvenation to the I/R-injured heart. Mechanistically, this selective macrophage response altered cardiac fibroblast activity and reduced border zone extracellular matrix (ECM) content and enhanced the mechanical properties of the injured area. The functional benefit of cardiac cell therapy is thus due to an acute inflammatory-based wound healing response that rejuvenates the mechanical properties of the infarcted area of the heart. Such results suggest a re-evaluation of strategies underlying cardiac cell therapy in current and planned human clinical trials.

---

Initial animal studies of adult stem or progenitor cell therapy for cardiac regeneration reported improved heart function with robust de novo myogenesis derived directly from the injected cells^10–13^. Many independent groups have since repeated these studies with an ever-increasing variety of adult stem cells and while functional improvement is reproducibly observed, nearly all have failed to observe new cardiomyocyte formation from these injected cells^14–17^. At the same time, clinical trials with adult stem cells in patients with acute myocardial infarction (MI) injury or decompensated heart failure have expanded worldwide over the past 17 years, with thousands of patients enrolled and over $1 billion USD in funding utilized^4,6^. However, while results of these trials have been disappointing, this might simply reflect ineffective trial design because the true mechanistic basis of cell therapy remains unknown^6^. For example, more recent literature has shown that the vast majority of transplanted adult stem or progenitor cells simply die within days of delivery into the hearts of ischemia-injured animal models^18,19^, and as such it remains unclear how they might function in a therapeutic manner, although a paracrine hypothesis has been proposed whereby injected cells temporarily release protective growth factors or RNA species^20–22^.

Over 15 types of adult stem cells show some level of efficacy in cardiac regenerative studies of ischemic injury in animal models^7,23^. Here we focused on 2 primary types: fractionated bone marrow mononuclear cells (MNCs), which were the earliest and most heavily used cell type in clinical trials^2^, and cardiac mesenchymal cells from the heart that express the receptor tyrosine kinase c-Kit, which have been termed cardiac progenitor cells (CPCs)^12,24^. As we will describe, our major finding was that injection of these cell-types induces a sterile immune response in the heart, so we also examined the effect of injecting zymosan, a non-cellular and potent activator of the innate immune response^25,26^. MNCs and CPCs were isolated from mice expressing a constitutive membrane TdTomato (mTomato) fluorescent reporter and zymosan was conjugated to Alexa Fluor 594 to allow tracking *in vivo* by red fluorescence. Isolated MNCs were a heterogeneous cell population consisting of all major hematopoietic lineages although monocytes and granulocytes were the most predominant (Extended Data Fig. 1a). CPCs expressed mesenchymal cell surface markers but were negative for markers of hematopoietic or endothelial cells (Extended Data Fig. 1b).

We first asked whether cell injection could activate endogenous regenerative programs in the murine heart in the absence of acute damage or disease, reasoning that this approach would allow us to isolate the specific biologic effects of injecting cells into the heart apart from the profound cellular complexity of an infarction injury model. Eight-week-old male and female *C57Bl/6J* mice received intra-cardiac injection of either strain-matched MNCs, 10 μg zymosan or saline. We chose a dose of 50,000 cells for these studies based on prior literature showing therapeutic benefit within this range^7,8^, while our zymosan dosing in adult mice was extrapolated from a recent report using zymosan injection into neonatal mouse hearts^27^. Injected material was delivered across 3 defined regions along the anterior wall of the left ventricle, mirroring the region of the heart most affected by MI (Fig. 1a). Importantly, none of the injected mTomato-labeled MNCs transdifferentiated into cardiomyocytes or endothelial cells (not shown). Distinct histological foci of acute local hyper-cellularity were observed with both cell and zymosan injection, as examined by confocal microscopy from heart sections 3 days or 2 weeks post-injection, suggestive of an inflammatory response (Fig. 1b). Indeed, activated CD68^+^ macrophages were significantly increased specifically within the area of injection at 3 days, with a diminishing effect by 2 weeks as the cells or zymosan were cleared (Fig. 1b, c). We also examined neutrophil levels by flow cytometry from dissociated hearts at 3 days, but we did not observe a significant increase with cell or zymosan injection (Extended Data Fig. 1c).

**Figure 1.**
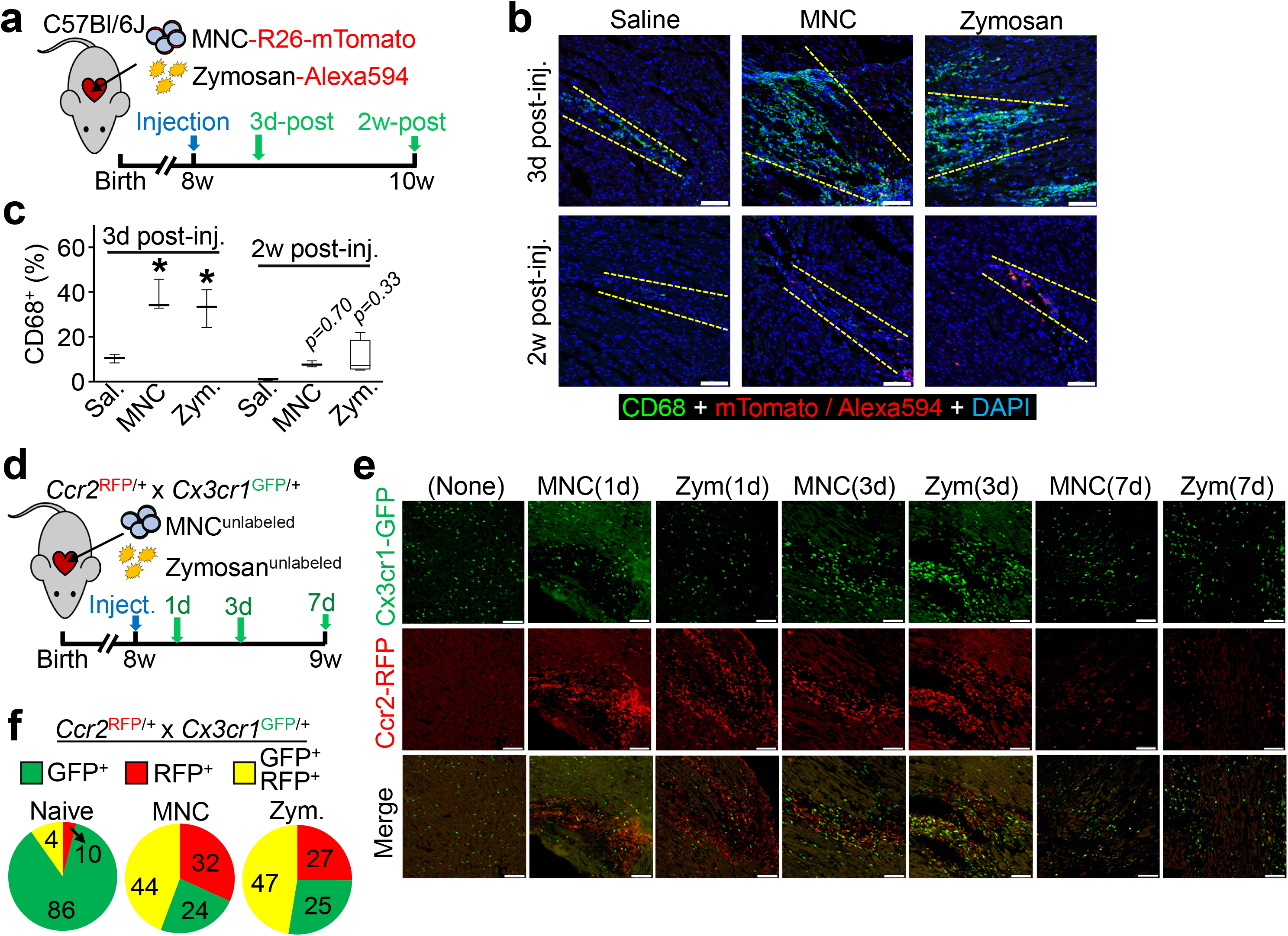
Cardiac cell injection causes local inflammation with accumulation of distinct macrophage subtypes. **a,** Experimental scheme used here in 8w-old male and female *C57Bl/6J* mice subjected to intra-cardiac injection of strain-matched bone marrow mononuclear cells (MNC), Alexa Fluor 594-conjugated zymosan (Zym.) or sterile saline (Sal.). Cells were isolated from *Rosa26*-mTomato mice on the *C57Bl/6* background. **b,** Representative confocal immunohistochemistry micrographs of hearts from *n*=4 mice (Zym; 2w-post) or *n*=3 mice (all other time points) showing activated CD68 macrophages (green) or the injected MNCs or zymosan (red). DAPI (blue) shows nuclei and dashed lines show injection sites. Data are from a minimum of 30 histological sections per mouse heart assessed from *n*=4 mice (Zym; 2w-post) or *n*=3 mice (all other time points). Scale bars = 100 μm. **c,** Quantitation of CD68^+^ cells as a percentage of total cells (DAPI^+^) imaged at areas of injection from the groups shown in (**b**). *p<0.05 versus saline by one-way ANOVA with Tukey’s post-hoc test. Data are summarized as box and whisker plots indicating the median value (black bar inside box), 25th and 75th percentiles (bottom and top of box, respectively), and minimum and maximum values (bottom and top whisker, respectively). **d,** Experimental scheme using 8w-old male and female *Ccr2*-RFP x *Cx3cr1*-GFP knock-in mice to simultaneously visualize CCR2^+^ and CX3CR1^+^ macrophage subsets in vivo after injection of MNCs or zymosan. Here MNCs were isolated from wild-type *C57Bl/6J* mice while zymosan was not conjugated to a fluorophore. **e,** Representative confocal micrographs from MNC or zymosan-injected hearts, versus naïve (uninjected) controls (minimum of 30 sections assessed per mouse heart from *n*=2 naïve control mice and *n*=3 MNC or *n*=3 zymosan-injected mice), showing endogenous RFP and GFP immunofluorescence from CCR2^+^ or CX3CR1^+^ macrophages, respectively, at the injection site over a 7-day time course. Scale bars = 100 μm. **f,** Distribution of CCR2^+^ and CX3CR1^+^ macrophage subtypes in hearts at 3 days post-injection. Pie charts reflect the proportion of RFP (CCR2^+^) or GFP (CX3CR1^+^) expressing macrophages, as well as CCR2^+^ CX3CR1^+^ double-positive (yellow) macrophages detected by flow cytometry, as a percent of total macrophages identified by staining for F4/80 and CD64. Data are from *n*=6 MNC and *n*=6 zymosan-injected mice and *n*=2 naïve (uninjected) mice.

To further profile the induction of macrophages with MNCs or zymosan injection, we employed a genetic approach in mice to differentially label CCR2^+^ and CX3CR1^+^ macrophages with either red (*Ccr2*-RFP^28^) or green (*Cx3cr1*-GFP^29^) fluorescence. CCR2 and CX3CR1 regulate the recruitment and/or activation of circulating monocyte-derived or tissue-resident macrophages, respectively^30–33^, and have been established as markers to broadly distinguish pro-inflammatory monocyte-derived macrophages (CCR2^+^) versus pro-healing macrophages (CX3CR1^+^ CCR2^-^)^34–39^. We delivered unlabeled MNCs or zymosan into 8-week-old *Ccr2*-RFP x *Cx3cr1*-GFP mice by intracardiac injection (Fig. 1d). Uninjured adult (non-injected) hearts showed GFP^+^ (CX3CR1^+^) tissue-resident macrophages throughout the myocardium while RFP^+^ (CCR2^+^) macrophages were largely absent (Fig. 1e). After 1 day, areas of MNC and zymosan injection in the heart showed a robust and highly localized influx of CCR2^+^ macrophages within the injection site while CX3CR1^+^ macrophages were largely restricted to the periphery of the injected area (Fig 1e). By 3 days these CX3CR1^+^ macrophages expanded within the injection area along with CCR2^+^ macrophages, but by 7 days levels of both macrophage types were reduced, indicative of a biphasic and temporal response. Flow cytometry analysis from these mice at 3 days also indicated a shift in overall macrophage subtype content from a largely CX3CR1^+^ population in the naïve state to a mix of CCR2^+^ and CCR2^+^ CX3CR1^+^ (double-positive) macrophages with either MNC or zymosan injection (Fig. 1f). Taken together these data suggest that the principal endogenous cellular response to intra-cardiac cell therapy or zymosan injection is an acute and localized inflammation through biphasic involvement of CCR2^+^ and CX3CR1^+^ macrophages that is largely cleared within 1 week.

One aspect of the proposed paracrine hypothesis of cell therapy introduced above is that the injected cells secrete effectors that cause endogenous cardiomyocytes to proliferate^20,21,40^, which we examined by immunohistochemistry from hearts injected with MNCs or zymosan (Fig. 2a). We used an antibody against PCM-1 to specifically mark cardiomyocyte nuclei^41^ and Ki67 to label nuclei with cell cycle activity (Fig. 2b). No appreciable increase in cardiomyocyte cell cycle activity was observed versus saline-injected controls, either at areas of injection or distally across the entire tissue (Fig. 2c). Another proposed effect of cell therapy is the activation of endogenous stem or progenitor cells, in particular cardiac-resident stem cells expressing c-Kit^21,40^. We and others^42–45^ have employed genetic lineage tracing of c-Kit^+^ cells *in vivo* and found that their endogenous contribution to cardiomyogenesis is negligible even after injury. Here we used tamoxifen-inducible *Kit*^MerCreMer/+^ x R-eGFP lineage tracing mice to directly examine new cardiomyocyte generation from endogenous c-Kit^+^ cells. Tamoxifen was administered over 6 weeks allowing for greater cumulative eGFP labeling. For these experiments we also injected cardiac progenitor cells (CPCs) that were isolated from the heart and selected for c-Kit positivity and then expanded in culture (CPCs), in addition to MNCs or zymosan (Fig. 2d). We observed very rare single c-Kit^+^-derived (eGFP^+^) cardiomyocytes in all treatment groups at a physiologically insignificant amount that was not greater with injected MNCs, CPCs, or zymosan versus saline (Fig. 2e, f). These data demonstrate that stem or progenitor cell injection does not cause endogenous cardiomyocyte proliferation or the induction of cardiomyocytes from endogenous c-Kit^+^ progenitor cells.

**Figure 2.**
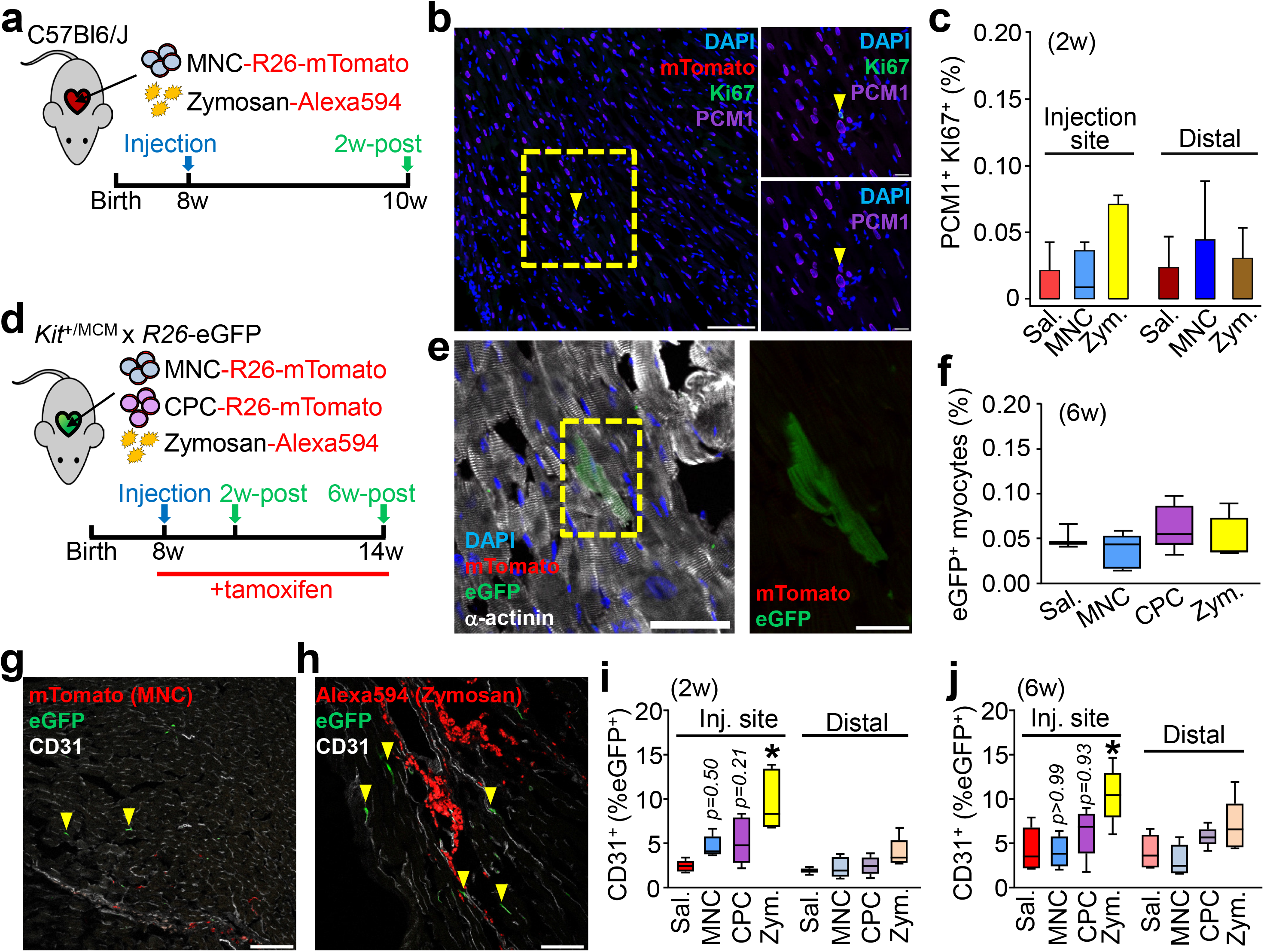
Cell or inflammatory therapy induces some endothelial cell but not cardiomyocyte formation. **a,** Schematic outline of experiments performed in panels **b** and **c** in 8w-old male and female *C57Bl/6J* mice with intra-cardiac injection of MNCs, zymosan, or saline and analyzed 2w later. **b,** Representative cardiac immunohistochemistry for Ki67 (green) and PCM1 (purple) from MNC-injected hearts. DAPI (blue) shows nuclei. Scale bar = 100 μM. A minimum of 45 histological sections were analyzed per mouse heart from *n*=4 MNC-treated mice or *n*=5 for all other groups of mice. Yellow box denotes area shown in higher magnification insets on the right. Yellow arrowhead denotes a cardiomyocyte with cell cycle activity. Scale bars for inset images = 10 μm. **c,** Quantitation of cardiomyocytes with cell cycle activity (PCM1^+^ Ki67^+^) as a percentage of all cardiomyocytes imaged (PCM1^+^) at 6w. Data are from a minimum of 45 histological sections analyzed per mouse heart from *n*=4 MNC-treated mice or *n*=5 for all other groups of mice. **d,** Schematic outline of experiments performed in panels (**e-j**) using c-Kit lineage tracing mice (*Kit*^*MerCreMer/+*^ *x R-eGFP*) injected with MNCs, CPCs, zymosan or saline, then analyzed 2w or 6w later. Tamoxifen was administered continuously (in chow) starting one day before cell injection. **e,** Representative cardiac immunohistochemistry at 6w for α-actinin (white) to show cardiomyocytes or eGFP (green) to show *Kit* allele-derived cells. DAPI (blue) shows nuclei. Scale bar = 50 μm. Yellow box highlights an eGFP^+^ cardiomyocyte, shown at greater detail to the right. Inset scale bar = 10 μm. **f,** Quantitation of percent *Kit* allele-derived eGFP^+^ cardiomyocytes relative to total cardiomyocytes counted. Data are from *n*=3 saline-treated mice or *n*=5 for all other groups of mice. **g, h,** Representative confocal cardiac immunohistochemistry images for CD31^+^ endothelial cells (white) and also showing MNCs (**g**) or zymosan (**h**) from injected hearts (red). Arrowheads denote CD31^+^ endothelial cells that are also eGFP^+^. Scale bars = 100 μm. **i, j,** Quantitation of percent eGFP^+^ endothelial cells relative to total endothelial cells counted, either 2w (**i**) or 6w (**j**) post-injection. Data are from *n*=6 Sal/2w or Zym/6w or *n*=5 all other groups of mice. *p<0.05 by one-way ANOVA with Tukey’s post-hoc test. All numerical data are summarized as box and whisker plots. Data and representative micrographs in **e-j** are from a minimum of 45 histological sections analyzed per individual mouse heart from the numbers of mice as indicated above.

We also assessed the formation of new endogenous endothelial cells from the c-Kit^+^ lineage in the hearts of *Kit*^MerCreMer/+^ x R-eGFP mice (Fig. 2g-j). Two weeks after injection, eGFP^+^ endothelial cells were significantly increased at the injection sites of zymosan-treated (Fig. 2h, 2i) hearts. In contrast we observed a non-significant increase with MNC or CPC injection (Fig. 2g, 2i) and by 6 weeks only zymosan injection still gave a significant increase in endothelial cells (Fig. 2j). However, zymosan persists the longest within the heart while CPCs and MNCs are essentially cleared by 2 weeks, (Fig. 2g, 2h). None of the treatments increased c-Kit^+^-derived endothelial cells in the distal areas of the heart, suggesting localized induction in areas of active inflammation. Thus, the acute inflammatory response associated with cell therapy or zymosan injection could have a mild cardioprotective effect through a transient increase in new capillary formation (see below).

We next investigated whether cell therapy or acute inflammation with zymosan could positively impact the function of the mouse heart following MI injury due to prolonged ischemia/reperfusion (I/R). We injected either 150,000 total strain-matched MNCs, CPCs, 20 μg zymosan or saline (doses increased to account for exacerbated cell loss in the injured heart) on each side of the heart’s infarct border zone in *C57Bl/6J* mice, 1 week post-I/R injury (Fig. 3a). Importantly, cell or zymosan injection into uninjured hearts did not alter LV structure or function (Extended Data Fig. 2 a-f). Injection of MNCs, CPCs or zymosan each significantly improved post-I/R cardiac function at 2w post-injection (3w post-I/R) as measured by fractional shortening (Fig. 3b), and this was associated with improvements in left ventricular end-systolic volume (LVESV; Extended Data Fig. 3a). In contrast, cell or zymosan therapy showed no change in end-diastolic volume (LVEDV; Extended Data Fig. 3b) or heart rate (Extended Data Fig. 3c) across any of the treatment groups at 2w post-therapy, suggesting no effect on ventricular remodeling, but instead that contractile properties of the heart were directly impacted. Importantly, the functional benefit persisted for at least 8w after injection, as indicated by increased fractional shortening in MNC or zymosan-treated mice (Fig. 3c). By 8w after injection, saline-treated I/R hearts also began to show ventricular dilation as indicated by a significant increase in LVEDV, but this was attenuated in the hearts of MNC-treated mice (data not shown).

**Figure 3.**
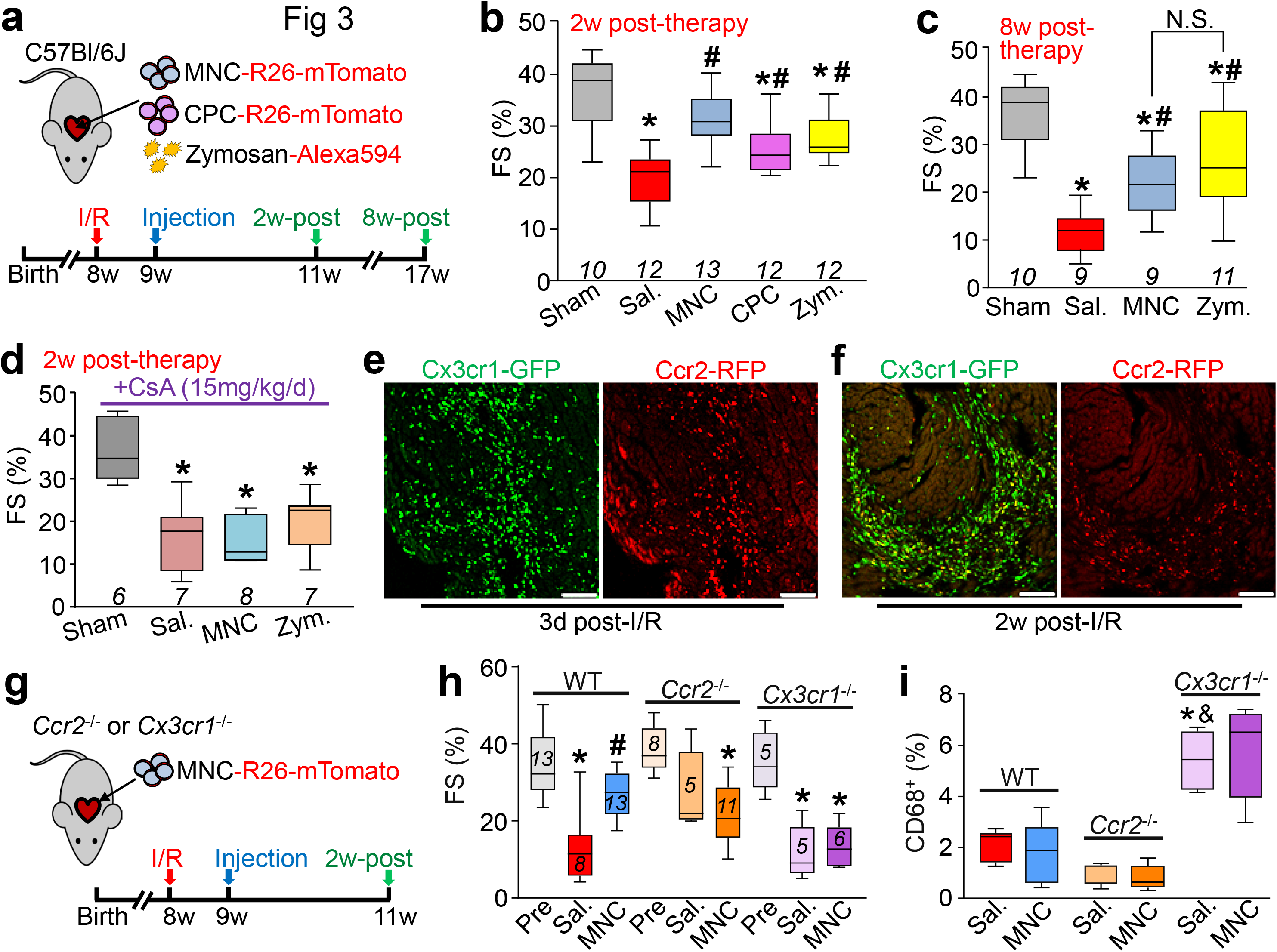
Cell or inflammatory therapy rejuvenates post-I/R heart function. **a,** Schematic outline of experiments performed in panels **b-c** in which 8w-old male and female *C57Bl/*6*J* mice received 120 min of myocardial ischemia followed by reperfusion (I/R) injury then intra-cardiac injection 1w later of MNCs, CPCs, zymosan, or sterile saline flanking the injury area, followed by analysis 2w or 8w later. **b-c,** Fractional shortening (FS) as measured by echocardiography in the groups indicated, 2w post-cell or zymosan therapy (**b**) or 8w post-cell or zymosan therapy (**c**). *p<0.05 vs Sham/Sal, or ^#^p<0.05 vs I/R/Sal. by one-way ANOVA with Dunnett’s post-hoc test. The same sham group is shown in **b** and **c** as these experiments were performed in parallel. **d,** FS as measured by echocardiography in male and female post-I/R mice that received cyclosporine A (CsA; 15 mg/kg body weight/d) delivered by osmotic minipump, starting one day before MNC or saline injection and continuing out for 2w post-injection. *p<0.05 vs Sham/Saline by one-way ANOVA with Dunnett’s post-hoc test. **e,** Confocal micrographs at the infarct border zone of hearts from male and female *Ccr2*-RFP x *Cx3cr1*-GFP knock-in mice (*n*=2 mice per group and timepoint with a minimum of 10 sections assessed per mouse heart) at either 3 days or 2 weeks post-I/R. **g,** Schematic outline of experiments performed in panels (**h**) and (**i**) in male and female *Ccr2*^-/-^ or *Cx3cr1*^-/-^ mice in the *C57Bl/6* background that were subjected to I/R then injected with MNCs or sterile saline 1w later. **h,** FS in *Ccr2*^-/-^ or *Cx3cr1*^-/-^ mice or wild-type *C57Bl/6* mice 3w post-I/R (2w post-cell injection). *p<0.05 vs Pre-I/R (Pre) or ^#^p<0.05 vs I/R/Sal. by one-way ANOVA with Tukey’s post-hoc test. **i,** Quantitation of CD68^+^ cells as a percentage of total cells (DAPI^+^) imaged at the infarct border zone, 3w post-I/R. *p<0.05 versus WT/Saline or ^&^p<0.05 versus *Ccr2*^-/-^/Sal. by one-way ANOVA with Tukey’s post-hoc test. The number (*n*) of mice in all experimental groups is indicated below or within the respective plot. All numerical data are summarized as box and whisker plots.

Our data indicated that the predominant effect of intracardiac cell therapy was localized and biphasic action of CCR2^+^ followed by CX3CR1^+^ macrophages, which was recapitulated with zymosan. To more specifically examine whether this inflammatory response was a mediator of post-I/R rejuvenation with cell therapy, we first treated mice with cyclosporine A (CsA), a broad-spectrum immunosuppressant, starting 1 day prior to cell injection. MNCs were used as they are the overwhelming cell-type used in human clinical trials^1,2^. Remarkably, CsA abrogated the restorative effects on cardiac function seen with MNC or zymosan injection after I/R injury (Fig. 3d), indicating that the immune response was required for the observed benefit. In addition, we injected freeze-thaw killed MNCs to address the paracrine hypothesis, and remarkably, dead cell debris similarly improved cardiac function post-I/R, further suggesting that the acute sterile immune response is a primary mechanism of action for cell therapy (Extended Data Fig. 3d).

Cardiac I/R injury itself is associated with a robust and temporally regulated recruitment of discrete myeloid cell populations^34,46,47^, which we also observed using our *Ccr2*-RFP x *Cx3cr1*-GFP mice. (Fig. 3e). Of note, this sequential expansion of CCR2^+^ followed by CX3CR1^+^ macrophages in the developing scar and infarct border zone was reminiscent of the localized pattern we observed with cell therapy or zymosan injection into naïve hearts, suggesting that cell therapy injection simply invokes another round of acute wound healing. To test this, we employed *Ccr2*^-/-^ or *Cx3cr1*^-/-^ gene-targeted mice. Although initial infarct sizes post-I/R were not different among *Ccr2*^-/-^ or *Cx3cr1*^-/-^ mice or strain-matched wild-type controls (not shown), *Ccr2* deficiency significantly improved cardiac function after I/R (Fig. 3h), consistent with previous reports^31,48,49^. Moreover, cell therapy by MNC injection in mice lacking *Ccr2* imparted no further functional benefit, given the already enhanced state of cardiac healing after I/R (Fig.3h). Loss of *Ccr2* showed a reduction in overall CD68^+^ cell content in the post-I/R heart with or without cell therapy, consistent with prior studies demonstrating that targeting circulating CCR2^+^ monocytes/macrophages can reduce inflammation in the heart^31,49^. By comparison, *Cx3cr1* null mice lacking tissue resident macrophage activity showed left ventricular dysfunction after I/R injury that was similar to wild-type controls, but these mice no longer benefitted from MNC therapy and showed a much greater total inflammatory response (Fig. 3h, i). This result suggests that endogenous tissue resident macrophages are necessary for a protective healing response associated with cell therapy. The increase in total inflammation in the I/R injured hearts of *Cx3cr1* null mice is consistent with recent data whereby these tissue-resident macrophages play an immunomodulatory role, dampening excess inflammation and promoting healing^34,50^.

Since cell therapy or zymosan injection produced only a transient increase in endothelial cell content in the heart it seemed unlikely to be the primary mechanism for protection. Thus, we hypothesized that a more fundamental mechanism was at play, such as an alteration in the properties of the ECM within the injury area and border zone. While overall infarct size in MNC-treated mice was unchanged versus controls (not shown), fibrotic burden specifically in the peri-infarct border zone was significantly decreased with MNC cell therapy (Fig. 4a, b). This was also observed with injection of non-viable MNCs, suggesting that it was primarily due to immunoreactivity and not active paracrine signaling (Fig. 4c). Tissue strips from the infarct region of saline or MNC-injected hearts were isolated and subjected to passive force-tension stress analysis. Remarkably, infarct strips from MNC-injected hearts produced a significantly greater change in passive force over increasing stretch (change in initial length [L_0_]; Fig. 4d). This was associated with a decrease in gene expression levels of several ECM and matricellular components in MNC versus saline treated hearts post-I/R (Fig. 4e). Together these data indicate that infarcts from MNC-injected hearts have improved mechanical properties and a reduction in fibrotic content versus saline controls. We also repeated the force-lengthening assay on infarct strips from post-I/R hearts injected with zymosan, which showed that these infarcts had an even larger improvement in passive force dynamics compared with either saline or MNC treatment (Extended Data Fig. 4a).

**Figure 4.**
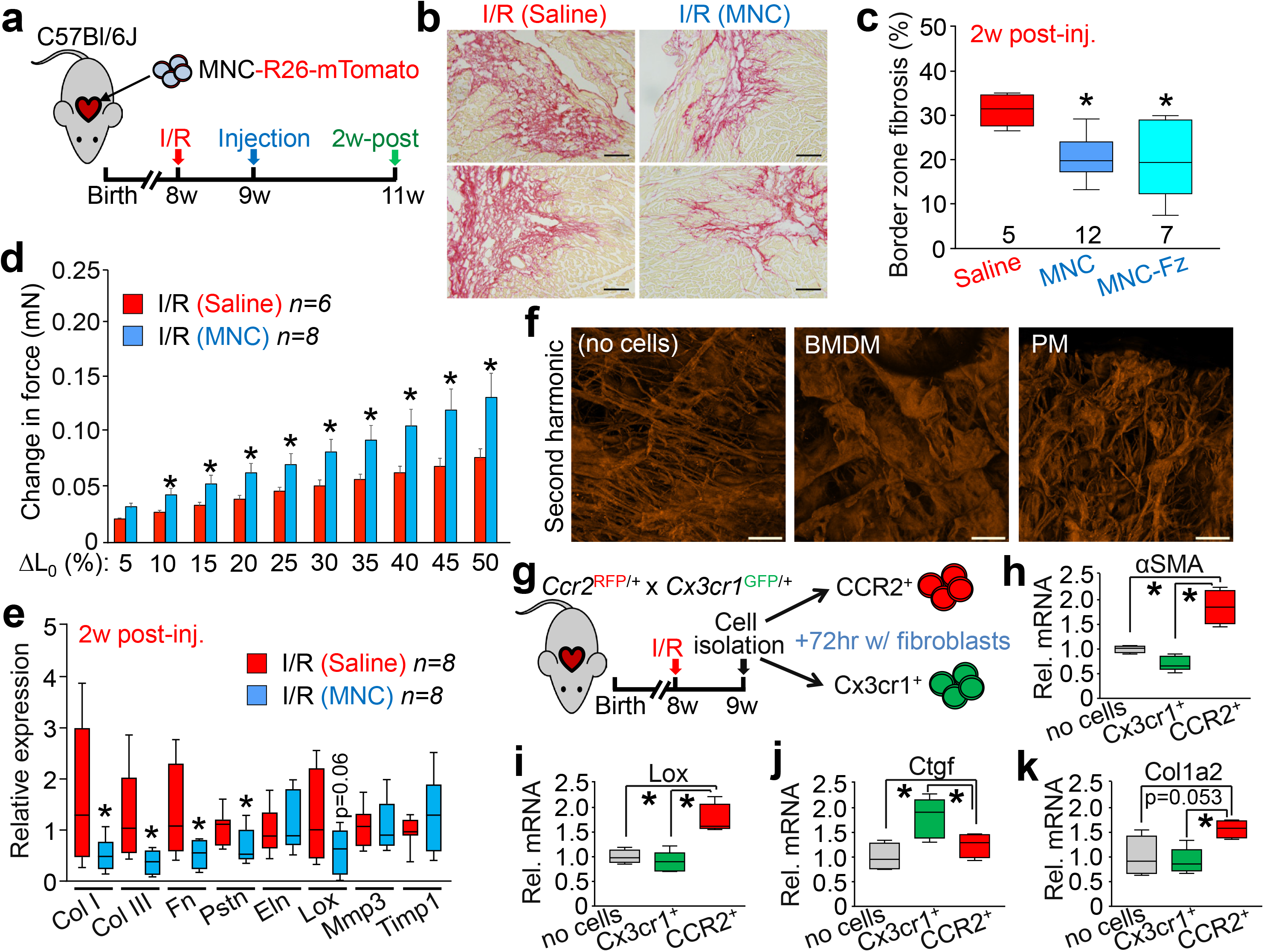
Cell therapy benefits the mechanical properties of the infarct via remodeling of the extracellular matrix. **a,** Schematic outline of experiments performed in panels **b-e** in this figure. **b,** Representative picrosirius red-stained cardiac histological images from the infarct border zone of 3w post-I/R mice subjected to MNC or saline injection. Fibrosis is shown in red. Scale bars = 100 μm. Images and quantitation in (**c**) are from *n*=5 saline-treated, *n*=12 MNC-treated, or *n*=7 freeze-killed MNC-treated mice, with a minimum of 20 histological sections assessed from each individual mouse heart. **c,** Quantitation of fibrotic area at the infarct border zone in MNC, freeze thaw-killed MNC, or saline-treated hearts (number of mice in each group shown below each box and whisker plot), 3w post-I/R. *p<0.05 versus I/R/Saline by one-way ANOVA with Tukey’s post-hoc test. **d,** Change in passive force generation over increasing stretch-lengthening (percent of L_0_) in isolated infarct strips from MNC or saline-treated hearts, 3w post-I/R. *p<0.05 versus I/R/Saline by Student’s 2-tailed t-test. **e,** Gene expression levels by RT-PCR for selected extracellular matrix (ECM) and matrix-associated genes in isolated infarct strips from MNC or saline-treated hearts, 3w post-I/R. *p<0.05 versus I/R/Saline by Student’s 2-tailed t-test. **f,** Representative confocal micrographs of pre-fabricated collagen patches that were seeded and cultured for 5 days with either bone marrow-derived macrophages (BMDM) or peritoneal macrophages (PM) isolated from wild-type male and female mice, versus cell-free control patches cultured in media. Fluorescence signal is from second harmonic generation microscopy using 840 nm light to allow for specific detection of native type I and II collagen. Scale bars = 100 μm. **g,** Schematic outline of experiments using activated cardiac macrophages isolated from post-I/R *Ccr2*-RFP x *Cx3cr1*-GFP knock-in mice using fluorescence activated cell sorting (FACS). CCR2^+^ and CX3CR1^+^ macrophages were then cultured with isolated cardiac fibroblasts for 72 hrs. **h-k,** Fibroblast mRNA was then used for RT-PCR to assess expression of smooth muscle α-actin (*Acta2* / αSMA, **h**), lysyl oxidase (*Lox*, **i**), connective tissue growth factor (*Ctgf*, **j**) or collagen 1 alpha 2 (*Col1a2*, **k**). Numerical data in (**d**) are presented as the mean + SEM from the number (*n*) of mice indicated in the figures. All other numerical data are summarized as box and whisker plots. Micrographs in **f** are representative of five different collagen patches seeded with cells pooled from *n*=4 mice (2 male and 2 female). Data in **g-k** are from five replicates generated over fibroblasts isolated from *n*=10 wild-type mice (6 male and 4 female) and macrophages isolated from *n*=6 *Ccr2*-RFP x *Cx3cr1*-GFP knock-in mice (3 male and 3 female).

To further define the mechanism, we performed a series of in vitro experiments using purified macrophages to determine how the acute induction of inflammation with cell therapy might impart structural and functional changes to the post-I/R region of the heart. We first isolated either bone marrow-derived (BMDM) or peritoneal (PM) macrophages from naïve mice, which have been used as broad models for circulating monocyte-derived or tissue-resident macrophages, respectively^51,52^. We cultured these macrophages on pre-fabricated collagen patches and then imaged the patches with second harmonic generation microscopy to examine collagen organization. Addition of macrophages to these patches, which lack fibroblasts or any other cell type, resulted in dramatic remodeling of the collagen matrix (Fig. 4f). This effect was highly distinct between BMDM, which produced wide folds throughout the patch, versus PM, which reorganized the matrix into thinner and interlaced sheets. To test whether a similar reorganization of collagen via macrophage activity was occurring in our cell or zymosan-treated hearts in vivo, we performed histological analysis on post-I/R hearts from *Ccr2*-RFP x *Cx3cr1*-GFP mice that received MNCs or zymosan, using a recently described collagen hybridizing peptide (CHP^53^) that specifically detects immature or denatured collagen (Extended Data Fig. 4b). Hearts from MNC or zymosan treated mice showed a CHP reactivity that was coincident with enhanced presence of CCR2^+^ and CX3CR1^+^ macrophages (Extended Data Fig. 4c), suggesting that sites of active inflammation within the areas of cell or zymosan injection were undergoing greater ECM remodeling. To address whether these activated macrophages could also be directly impacting cardiac fibroblasts, we purified CCR2^+^ or CX3CR1^+^ macrophages from hearts at 7 days post-I/R (Fig. 4g) and cultured them for 72 h with freshly isolated cardiac fibroblasts. Gene expression analysis of these fibroblasts by RT-PCR revealed that CCR2^+^ macrophages imparted an activating signal, as indicated by increased fibroblast expression of smooth muscle α-actin (*Acta2* / αSMA), lysyl oxidase (*Lox*), and collagen 1 alpha 2 (*Col1a2*) (Fig. 4h, i, k). In contrast, CX3CR1^+^ macrophages slightly reduced expression of these genes but increased fibroblast expression of connective tissue growth factor (*Ctgf*) (Fig. 4j), which has been associated with maintenance of the ECM as well as angiogenesis^54^. Taken together, these results demonstrate that acute localized inflammatory stimuli from CCR2^+^ and tissue-resident CX3CR1^+^ macrophages due to cell therapy improves the passive mechanical properties of the injured area by influencing the activity of cardiac fibroblasts.

Two of the most highly used adult stem cells, MNCs and CPCs, improved cardiac function in mice when delivered directly into both lateral sides of the infarct border zone, in agreement with previous data^10,13^. However, the same improvement was obtained with zymosan, a member of the pathogen-associated molecular pattern (PAMP) family that primarily acts via toll-like receptors to induce acute inflammation^26^. The observed functional benefit required macrophage-mediated inflammation, as shown using genetic or pharmacological inhibition. Most progenitor cells injected into the heart rapidly die^7,8,18,19^, and dying cells release damage-associated molecular patterns (DAMPs), cellular fragments that like PAMPs, are highly immunogenic^26,55,56^. Indeed, injection of freeze-thawed killed MNCs, containing DAMPs but incapable of active paracrine factor production, imparted equivalent benefit as live cells. However, our results do not rule out an additive effect of paracrine factors or miRNAs released over time from living injected cells.

The transitory and localized nature of this inflammatory response is likely crucial in producing a benefit, as excessive and chronic inflammation throughout the heart appears to be universally pathological^56,57^. In this context the very low retention and rapid clearance of injected adult stem cells may actually be advantageous. These observations may also explain why direct injection into the parenchyma of the heart, as done in the majority of animal studies^7–9^, is universally efficacious while pre-clinical studies using systemic vascular infusion of progenitor cells as performed in the majority of human clinical trials have shown mixed results^4,6,58,59^. Indeed, it is uncertain how systemic vascular delivery of progenitor cells could provide benefit to the heart given the mechanism of action we proposed here, as the vast majority of infused cells do not take up residence in the heart nor persist in the circulatory system^60,61^.

Our study was designed to examine the mechanistic basis of adult cardiac stem cell therapy given the new consensus that direct cardiomyocyte regeneration is no longer a tenable mechanism^17,62^ and that the adult mammalian heart is largely non-regenerative and does not contain a resident stem cell population^63–65^. As suggested 10 years ago^66^, we observed that the acute inflammatory response is the overwhelming mechanism of benefit behind cell therapy to the post-MI injured heart. Previous studies using severe genetically immunodeficient animals have also demonstrated that the benefit of injecting a type of adult bone marrow cell in the heart post-MI was lost^67^. We identified a unique mechanism whereby stimulating the intrinsic wound healing cascade and sequential activity of circulating CCR2^+^ followed by CX3CR1^+^ tissue resident macrophages positively impacted the ECM around and within the infarcted region of the heart, such that functional performance was significantly improved. This acute inflammatory response provides instruction to resident fibroblasts in modifying the border zone ECM and its mechanical properties^46,68–70^. The dynamics and cell-specific regulation of inflammation by macrophage subsets in the heart is a growing area of study^35,36,50,70^, such that the timing and rapid clearance of immune cells is also key to proper healing^71^. Hence, it might be warranted to re-evaluate current and planned cell therapy-based clinical trials to maximize the effects of the most prevalent underlying biologic mechanism of action demonstrated here.

## METHODS SUMMARY (also see supplementary methods)

Candidate cellular therapeutics (MNCs or CPCs) were isolated from mice with a genetically-encoded fluorescent mTomato reporter and delivered by intra-cardiac injection, either at baseline or after I/R injury, into strain-matched animals that were either wild-type, genetically null for macrophage subtypes, or carrying *Kit*-MerCreMer and Rosa26-eGFP transgenes (R26-eGFP, for genetic lineage tracing). Cardiac function, inflammation, fibrosis, and cellular regeneration were assessed by echocardiography, immunohistochemistry, or flow cytometry. See Supplementary Methods for more detailed information and Supplementary Tables 1 and 2 for description of primary antibodies and RTPCR primers used.

## Supplementary Information

Extended Data Figure 1

Extended Data Figure 2

Extended Data Figure 3

Extended Data Figure 4

Supplementary Table 1

Supplementary Table 2

Supplementary Methods

## Acknowledgments

This work was supported by grants from the National Institutes of Health to J.D.M., S.S., and M.N. J.D.M. was also supported by the Howard Hughes Medical Institute. R.J.V. was supported by a National Research Service Award from the NIH (F32 HL128083). All flow cytometric data were acquired using equipment maintained by the Research Flow Cytometry Core in the Division of Rheumatology at Cincinnati Children’s Hospital Medical Center.

## Author Contributions

J.D.M. and R.J.V. conceived the study. R.J.V., M.M., M.A.S., H.K., A.K.J., J.A.S., A.J.Y. and V.H. performed experiments and generated all the data shown in the manuscript. S.S. provided oversight and technical help along with J.A.S. in measuring myocardial scar mechanical properties. M.N. provided theoretical assessment of the project and advice in experimental design.

## Conflict of Interest or Competing Financial Interest

None

