## Supplemental figures, Tables, Methods for "An acute immune response underlies the benefit of cardiac adult stem cell therapy"

Extended Data Figure 1

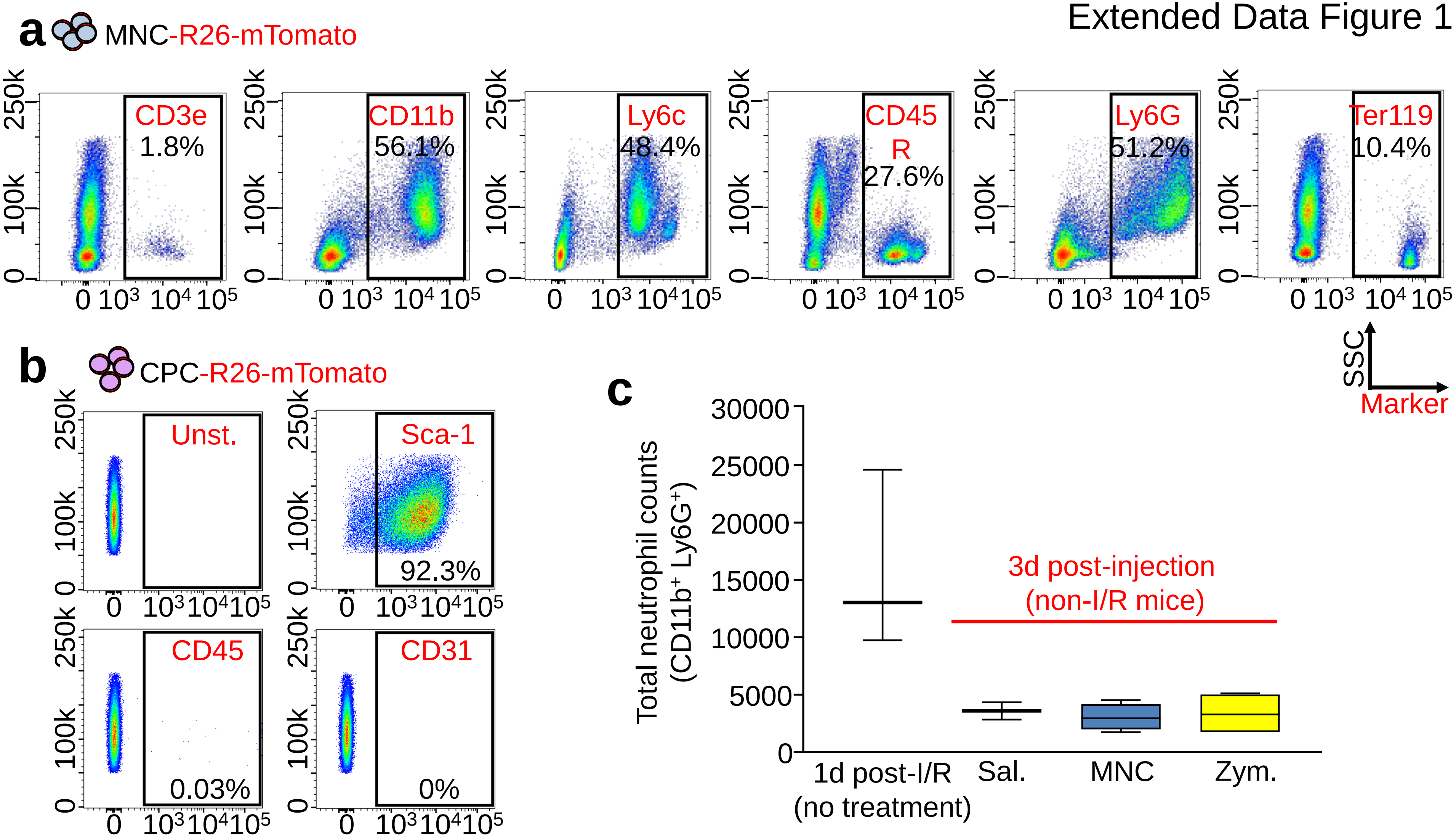

Extended Data Figure 2

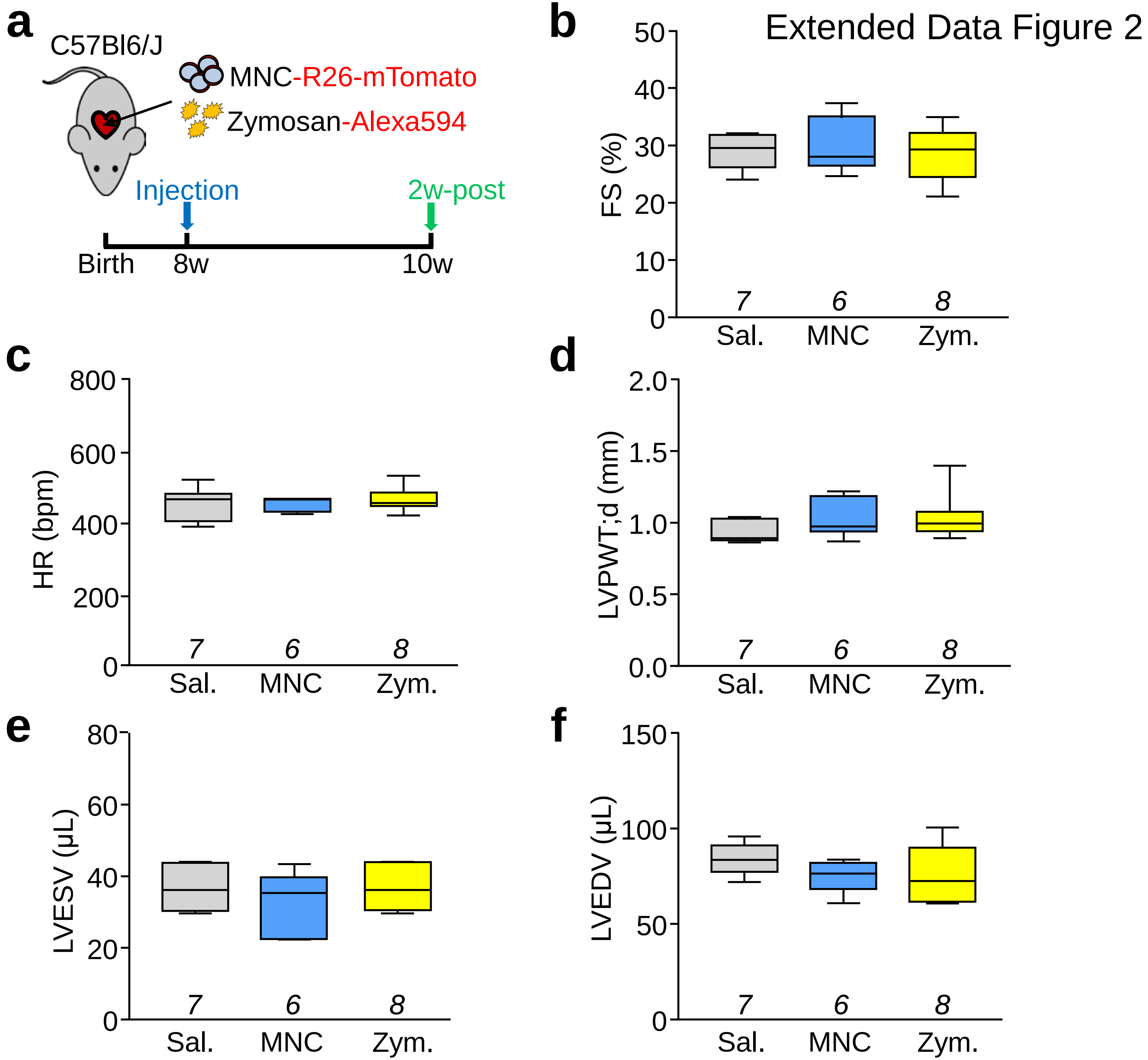

Extended Data Figure 3

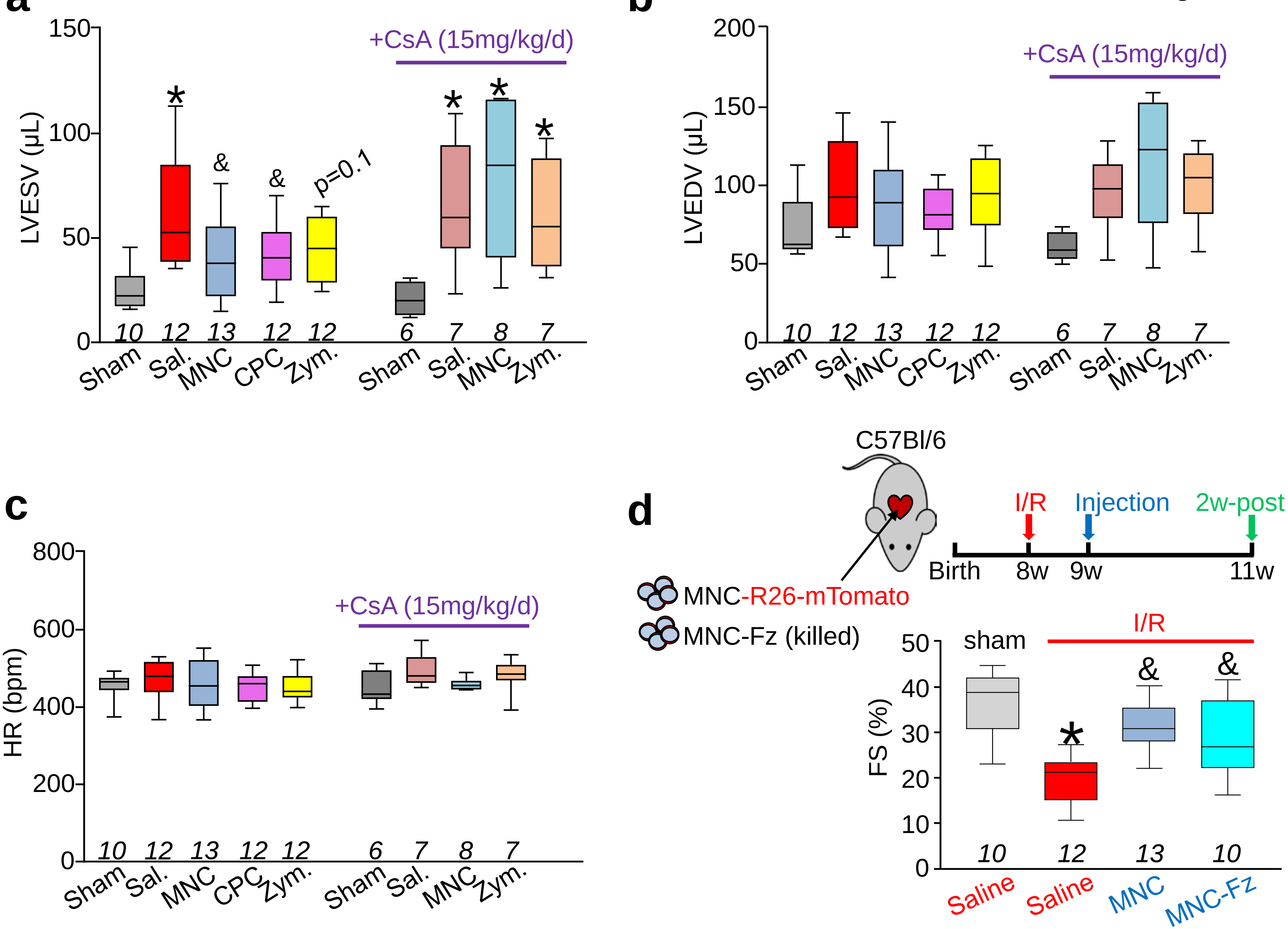

Extended Data Figure 4

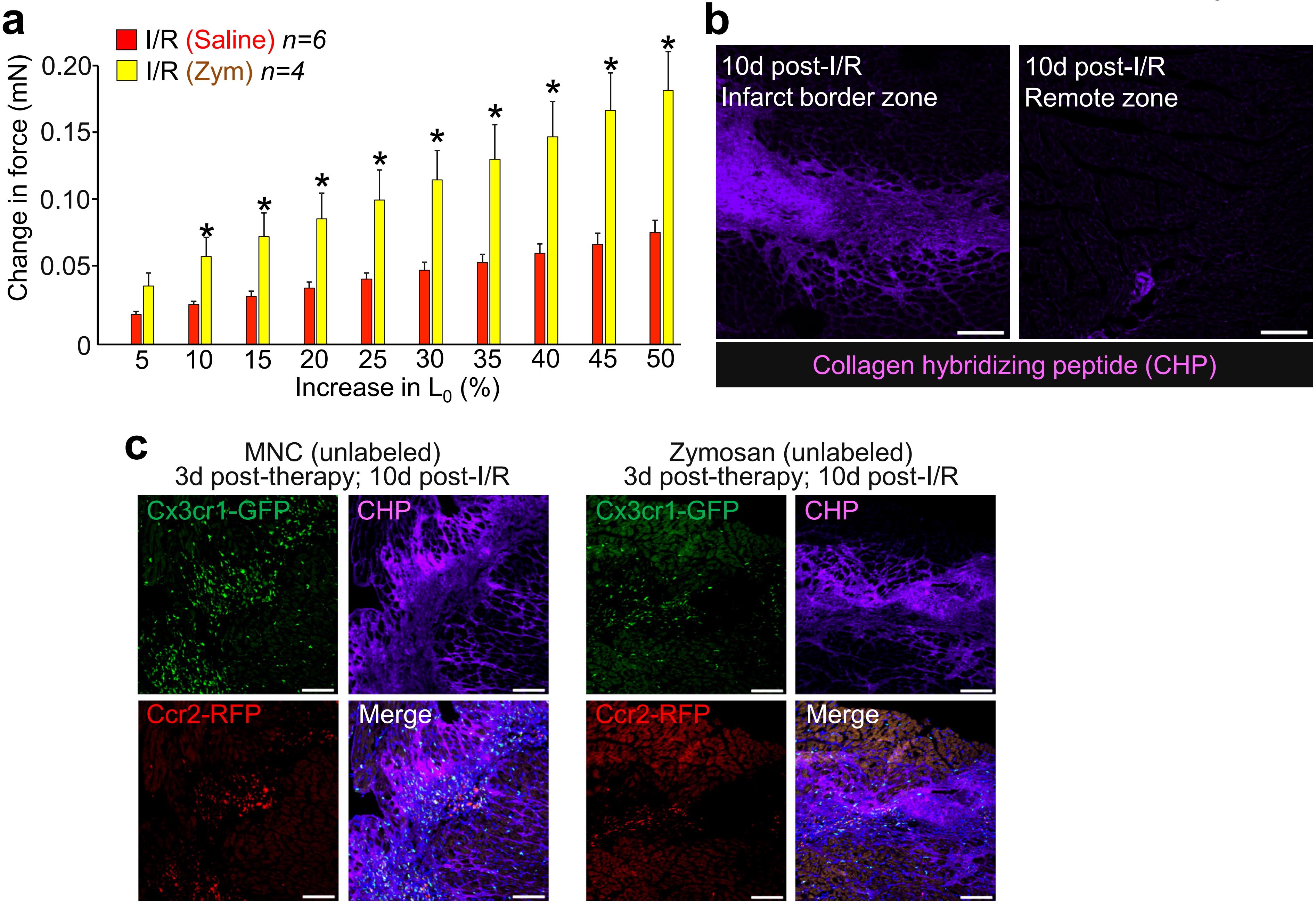

### EXTENDED DATA FIGURE LEGENDS

**Extended Data Figure 1 │ Characterization of cells used in the injection studies and initial neutrophil response to injections. a,** Flow cytometry analysis of MNCs isolated for intra-cardiac injection. Singlet cells were selected by forward and side scatter properties followed by mTomato-positivity. **b**, Flow cytometry plots for CPCs immunolabeled with antibodies against mesenchymal, endothelial, or hematopoietic lineages as indicated in the plots. An unstained negative control (Unst.) plot is also shown. Gating was determined versus unstained negative controls. **c,** Quantitation via flow cytometry of total neutrophil levels in dissociated hearts from MNC, zymosan, or saline-injected male and female *C57Bl/6* mice, 3 days post-injection. As a comparison, data from *C57Bl/6J* mice isolated 1 day after I/R injury are also shown. Data are from *n*=4 (MNC, Zym.) or *n*=2 (Sal. 1d post-I/R, mice). Numerical data are summarized as box and whisker plots indicating the median value (black bar inside box), 25th and 75th percentiles (bottom and top of box, respectively), and minimum and maximum values (bottom and top whisker, respectively).

**Extended Data Figure 2 │ Basal cardiac structure and function following cell or zymosan injection. a,** Schematic outline of all experiments performed in this figure in which 8w-old male and female *C57Bl/6J* mice received intra-cardiac injection of MNCs, zymosan (Zym.) or sterile saline (Sal.) and were assessed by echocardiography after 2w. **b,** Echocardiography measured fractional shortening percentage (FS%); **c**, heart rate (HR) as beats per minute (bpm) under isoflurane anesthesia; **d**, left ventricular posterior wall thickness in diastole (LVPWT;d) in millimeters; **e**, left ventricular end-systolic volume (LVESV) in microliters; **f**, and left ventricular end-diastolic volume (LVEDV) in microliters. All values in **b-f** were unchanged with injection of MNCs or zymosan versus saline. The number (*n*) of mice for each group is indicated below the respective plot. All numerical data are summarized as box and whisker plots indicating the median value (black bar inside box), 25th and 75th percentiles (bottom and top of box, respectively), and minimum and maximum values (bottom and top whisker, respectively).

**Extended Data Figure 3 │ Additional echocardiographic parameters and effect of freeze-thaw killed MNCs post-I/R injury. a,** Echocardiography to measure left ventricular end-systolic volume (LVESV) in microliters; **b**, left ventricular end-diastolic volume (LVEDV) in microliters; **c**, or heart rate under isoflurane anesthesia in mice that received intra-cardiac injection of MNCs CPCs, zymosan, or sterile saline, 3w post-I/R. $^{*}p<0.05$ vs Sham/Saline or $^{\&}p<0.05$ vs I/R/Saline by one-way ANOVA with Dunnett's post-hoc test. **d,** Schematic in mice in which mTomato-labeled MNCs or freeze-thaw (Fz)-killed MNCs were injected 1w after I/R and then 2w later cardiac ventricular fractional shortening (FS%) percentage was measured by echocardiography. $^{*}p<0.05$ vs Sham/Saline or $^{\&}p<0.05$ vs I/R/Saline by one-way ANOVA with Dunnett's post-hoc test. The sham, I/R+Saline, and I/R+MNC groups shown here in (**d**) are the same data as also shown in Main Fig. 3 as these studies were performed in parallel. All numerical data are summarized as box and whisker plots indicating the median value (black bar inside box), 25th and 75th percentiles (bottom and top of box, respectively), and minimum and maximum values (bottom and top whisker, respectively). The number (*n*) of mice for each group is indicated below the respective plot.

**Extended Data Figure 4 │ Mechanical and structural improvements in cell therapy or zymosan-treated hearts post-I/R. a,** Change in passive force generation over increasing stretch-lengthening (percent of $L_0$) in isolated infarct strips from zymosan or saline-injected hearts, analyzed 3w after surgical injury (injection of zymosan or saline occurred 2w before harvesting mice). $^{*}p<0.05$ versus I/R/Saline by Student's two-tailed t-test. The I/R+Saline data shown here are the same as in Main Fig. 4 as these studies were performed in parallel. Numerical data are presented as the mean + SEM from the number (*n*) of mice indicated in the figures. **b,** Representative confocal micrographs from heart histological sections from *C57Bl/6* mice at 10 days post-I/R showing the infarct border zone versus remote myocardium and labeled with a biotin-conjugated collagen hybridizing peptide (CHP) that detects immature or denatured collagen. CHP labeling was detected with a streptavidin-conjugated Alexa647 secondary antibody (purple). **c,** Representative confocal micrographs of heart sections from the post-I/R border zone of *Ccr2*-RFP x *Cx3cr1*-GFP mice that received intra-cardiac injection of MNCs or zymosan at 7d post-I/R and were analyzed after an additional 3d. Endogenous RFP (red) or GFP (green) fluorescence shows $CCR2^+$ or $CX3CR1^+$ macrophages, respectively. Sections were treated with CHP (purple) as in (**b**) to visualize immature collagen. Scale bars = 100 µm.

**Supplementary Table 1 │ Antibodies and antibody dilutions used in this study**

**Supplementary Table 2 │ Primers used for all RT-PCR analyses in this study**

**Supplementary Table 1 │ Antibodies and Antibody Dilutions Used in this Study**

| Reagent | Manufacturer | Catalog # | Isotype | Dilution (cryosections) | Dilution Flow cytometry) |
|---|---|---|---|---|---|
| CD68 antibody [FA-11] | Abcam | ab53444 | Rat monoclonal | 1:100 | |
| Brilliant Violet 421™ anti-mouse CD64 (FcγRI) | BioLegend | 139309 | Mouse IgG1, κ | | 1:100 |
| APC anti-mouse F4/80 [BM8] | BioLegend | 123116 | Rat IgG2a, κ | | 1:100 |
| Ki-67 Monoclonal Antibody (SolA15) | Thermo Fisher | 14-5698-82 | Rat IgG2a, κ | 1:100 | |
| PCM1 Antibody | Novus Biologicals | NBP1-87196 | Rabbit polyclonal | 1:100 (with antigen retrieval) | |
| Monoclonal Anti-α-Actinin (Sarcomeric) antibody produced in mouse | Sigma-Aldrich | A7811 | clone EA-53, ascites fluid | 1:100 | |
| Purified Rat Anti-Mouse CD31 Clone MEC 13.3 | BD Pharmigen | 553370 | Rat Lewis IgG2a, κ | 1:100 | |
| Collagen Hybridizing Peptide, Biotin Conjugate (B-CHP) | 3Helix | BIO300 | | 200μL of 15μM stock per section | |
| Biotin anti-mouse CD3 (145-2C11) | eBioscience | 88-7774-75 | | | 1:50 |
| Biotin anti-mouse CD11b (M1/70) | eBioscience | 88-7774-75 | | | 1:100 |
| Biotin anti-mouse CD45R/B220 (RA3-6B2) | eBioscience | 88-7774-75 | | | 1:100 |
| Biotin anti-mouse Ly-6G (RB6-8C5) | eBioscience | 88-7774-75 | | | 1:100 |
| Biotin anti-mouse Erythroid marker (TER-119) | eBioscience | 88-7774-75 | | | 1:100 |
| Pacific Blue™ anti-mouse Ly-6A/E (Sca-1) | BioLegend | 108120 | | | 1:100 |
| CD45-APC Clone 30-F11 | BD Pharmigen | 559864 | | | 1:100 |
| Anti-Mouse CD31 (PECAM-1) eFluor® 450 | eBioscience | 48-0311-82 | | | 1:100 |

**Supplementary Table 2 │ Primers Used for RT-PCR Analysis in this Study**

| Transcript | Forward Primer | Reverse Primer | Experiment |
|---|---|---|---|
| Collagen 1a1 | 5’- AATGGCACGGCTGTGTGCGA | 5’- AACGGGTCCCCTTGGGCCTT | Figure 4e |
| Collagen 3a1 | 5’- TCCCCTGGAATCTGTGAATC | 5’- TGAGTCGAATTGGGGAGAAT | Figure 4e |
| Fibronectin | 5’AAGGCTGGATGATGGTGGAC | 5’TGAAGCAGGTTTCCTCGGTTG | Figure 4e |
| Periostin | 5’- ACGGAGCTCAGGGCTGAAGATG | 5’- GTTTGGGCCCTGATCCCGAC | Figure 4e |
| Elastin | 5’CAGCTAAATACGGTGCTGCTG | 5’AATCCGAAGCCAGGTCTTG | Figure 4e |
| Lox | 5’TGATGCCAACACCCAGAGGA | 5’CGAATGTCACAGCGCACAAC | Figure 4e, 4i |
| Mmp3 | 5’CTTCTGCAACTCCGACATCGT | 5’GGGGCATCTTACTGAAGCCTC | Figure 4e |
| Timp1 | 5’GCAACTCGGACCTGGTCATAA | 5’CGGCCCGTGATGAGAAACT | Figure 4e |
| Gapdh | 5’TTGTCTCCTGCGACTTCAAC | 5’GTCATACCAGGAAATGAGCTTG | Figure 4e |
| Acta2 | 5'CTTCGTGACTACTGCCGAGC | 5'AGGTGGTTTCGTGGATGCC | Figure 4h |
| Ctgf | 5'AGAACTGTGTACGGAGCGTG | 5'GTGCACCATCTTTGGCAGTG | Figure 4j |
| Collagen 1a2 | 5'GGCCCCCTGGTATGACTGGCT | 5'CGCCACGGGGACCACGAATC | Figure 4k |
| Gapdh | 5'TGTCGTGGAGTCTACTGGTG | 5'ACACCCATCACAAACATGG | Figure 4h-k |

**Supplementary Methods:**

### Mice

This study was performed entirely in mice, using transgenic mouse models either commercially available or generated as described below. No human subjects or human material was used. All experiments involving mice were approved by the Institutional Animal Care and Use Committee (IACUC) at Cincinnati Children's Hospital under protocols IACUC2015-0047 and IACUC2016-0069. All procedures were performed in compliance with institutional and governmental regulations under PHS Animal Welfare Assurance number D16-00068 (A3108-01). The generation and characterization of mice carrying the tamoxifen-inducible MerCreMer recombinase cDNA within the *Kit* allele (*Kit*$^{MerCreMer/+}$) and reporter mice carrying the Cre-regulated LoxP-stop cassette and eGFP within the *Rosa26* gene locus, *Rosa26-eGFP* (R-GFP) were previously described[42]. All other mouse strains used were purchased from Jackson Labs, as follows: *C57Bl/6J*; (#000664), constitutive mTomato expressing mice targeted in the *Rosa26* locus for bone marrow mononuclear cell (MNC) or cardiac progenitor cell (CPC) isolation; (*B6.129(Cg)-Gt(ROSA)26Sortm4(ACTB-tdTomato,-EGFP)Luo/J*, #007676), *Ccr2* gene-deleted mice; (*B6.129S4-Ccr2tm1Ifc/J*, # 004999), *Cx3cr1* GFP knock-in mice (Cx3cr1 homozygotes are nulls); *B6.129P-Cx3cr1tm1Litt/J*, #005582), *Ccr2* RFP knock-in mice; (*B6.129(Cg)-Ccr2tm2.1Ifc/J #017586).* Both male and female sexes were used in all experiments, at age ranges indicated in the figures and main text for each experiment. Mice were housed single-sex at a maximum of 4 animals per cage in a specific pathogen free (SPF), temperature-controlled vivarium under a 12 hr light/dark cycle with ad libitum access to food and water.

### Preparation of Cell or Inflammatory Therapies

To generate MNCs for injection, whole bone marrow was first isolated by flushing dissected femurs and tibiae of 10-12w-old *Rosa26*-mTomato expressing mice or *C57Bl/6J*; mice with 10 mL of sterile Hanks Balanced Salt Solution (HBSS, Fisher Scientific #SH3058801) + 2% bovine growth serum (BGS, Fisher Scientific, #SH3054103) + 2 mM EDTA as previously described[72]. This suspension was then filtered through a 40 µM mesh strainer (Fisher Scientific #22-363-547), centrifuged at 400 g for 10 min at 4 °C, resuspended in 3 mL of sterile saline, and layered on top of 4 mL of Ficoll Paque Plus (GE Healthcare #17-1440-02). Cells were then centrifuged at 2500 g for 30 min at 4 °C in a swinging bucket rotor centrifuge without brakes. MNCs were isolated by removal of the resulting thin mononuclear cell layer (second layer from the top). Total MNCs were counted with a hemocytometer, washed twice with sterile saline, and resuspended in sterile saline at a final concentration of either $2.5x10^6$c/mL (for injection into uninjured hearts, final dose 50000 cells) or $7.5x10^6$c/mL (for injection into post-I/R hearts, final dose 150000 cells). The full intra-cardiac injection procedure is described in the following section (Mouse Procedures). Cell viability was tested by incubating an aliquot of the MNC suspension with eFluor 450 Fixable Viability Dye (eBioscience #65-0863-18) and found to be over 90% viable at the time of injection. All MNC preparations for injection were combined from an equal number of male and female mice. For experiments that used non-viable MNCs (freeze-thawed; Fz), this final MNC suspension was split into two equal aliquots. One aliquot was placed immediately at -80 °C for 10 min, followed immediately by incubation at +55 °C for 10 min, and this was repeated for a total of 3 freeze-thaw cycles.

To generate CPCs for injection, hearts from 10-12w-old *Rosa26*-mTomato expressing male and female mice were rapidly excised and briefly rinsed in cold 1X PBS. Single-cell suspensions from these hearts were prepared according to previously published protocols[73,74] with minor modifications, as follows. The atria were removed, and the ventricles were minced on ice using surgical scissors into approximately 2 mm pieces (8-10 pieces per mouse heart). Each dissociated ventricle was transferred into 2 mL of digestion buffer in 1 well of a 12-well tissue culture plate. Digestion buffer consisted of 2 mg/mL collagenase type IV (Worthington, #LS004188), 1.2 U/mL dispase II (Roche, #10165859001) and 0.9 mM $CaCl_2$, in 1x HBSS. Tissues were incubated at 37°C for 20 min with gentle rotation followed by manual trituration 12-15 times with a 10 mL serological pipette, such that all of the tissue pieces were able to pass through the pipette. The tissues were settled by sedimentation and the supernatant was passed through a 40 μM mesh strainer and stored on ice. Two milliliters of fresh digestion buffer was added, followed by 2 additional rounds of incubation, trituration and replacement of supernatant with fresh digestion buffer, except trituration was performed with a 5 mL serological pipette for round 2 and a 1 mL p1000 pipette tip (USA Scientific, #1112-1720) for round 3. The pooled supernatant from the 3 rounds of digestion was washed with sterile PBS and centrifuged at 200 g for 20 min at 4 °C in a swinging bucket rotor centrifuge without brakes. The pellet was resuspended in flow cytometry sorting buffer, consisting of 1x HBSS supplemented with 2% bovine growth serum (BGS) and 2 mM EDTA, and incubated with anti-c-Kit microbeads (Miltenyi Biotec #130-091-224) for 20 min at 4°C with gentle rotation. The suspensions were washed twice with sorting buffer and c-Kit$^+$ cells were enriched via positive selection over Miltenyi Biotec LS columns (#130-042-401) using benchtop magnetic cell separation (MACS) according to manufacturer's instructions. Cells isolated via positive MACS selection for c-Kit were cultured in DMEM/F12/Glutamax media (Gibco #10565-018) supplemented with 10% fetal bovine serum (Sigma, #F2442), 0.2% insulin/transferrin/selenium (ITS, Lonza, #17-838Z), 20 μg/mL basic recombinant human fibroblast growth factor (Promega, #G5071), $10^3$ U/mL leukemia inhibitor factor (Millipore, #ESG1106), 20 μg/mL epidermal growth factor (Sigma, #E9644) and 1% penicillin-streptomycin (Fisher Scientific, #30-002-CI). These c-Kit isolated cells from the heart are referred to as CPCs and they were expanded in culture for 12-15 passages before being used for injection, at which point cells were washed 3 times with sterile saline, trypsinized, counted, and resuspended in sterile saline at a final concentration of either $2.5 \times 10^6$c/mL (for injection into uninjured hearts, final dose 50000 cells) or $7.5 \times 10^6$c/mL (for injection into post-I/R hearts, final dose 150000 cells). As with MNCs, suspensions consisted of pooled CPCs from an equal number of male and female mice.

Alexa Fluor-594 conjugated zymosan, or unconjugated zymosan, were purchased from Thermo Fisher (zymosan A [S. cerevisiae] BioParticles, Alexa Fluor 594 conjugate, #Z-23374, zymosan A [S. cerevisiae] BioParticles, unlabeled, # Z2849). A suspension of either 1 mg/mL (for injection into uninjured hearts, final dose 10 μg) or 2 mg/mL (for injection into post-I/R hearts, final dose 20 μg) was prepared for injection in sterile saline according to manufacturer's instructions.

**Mouse Procedures**

To deliver cell or inflammatory therapies by intra-cardiac injection, mice were anesthetized using isoflurane inhalation (to effect), intubated, and a left lateral thoracotomy was performed. A 25 μL gas-tight syringe (Hamilton, #7654-01) fitted with a 33-gauge needle (Hamilton, #7803-05) was used for injections. For experiments in mice without injury, 20 μL of MNCs at a concentration of $2.5 \times 10^6$c/mL (50000 cells total) was injected over 3 regions of the LV (6.7 μL per injection). For experiments with zymosan, 10 μL of a 1 mg/mL suspension was injected over 3 regions of the LV (3.3 μL per injection, 10 μg zymosan total). Injections were equidistant along the anterior wall of the left ventricle (LV), with the needle entering just parallel to the long axis of the ventricle to avoid entering the LV chamber. For experiments with injections occurring after cardiac injury, 2 injections were performed, one on either side of the infarct zone, as follows. Twenty microliters total of MNCs or CPCs at a concentration of $7.5 \times 10^6$c/mL (150,000 cells total) was injected (10 μL per injection). For experiments with zymosan, 10 μL total of a 2 mg/mL suspension was injected (5 μL per injection, 20 μg zymosan total).

To induce cardiac injury, we used a modified surgical model of ischemia with reperfusion (I/R) via temporary left coronary artery ligation as previously described[75], except here we waited 120 min before inducing reperfusion so that more complete killing of the ischemic zone occurred to enhance reproducibility. After each surgical procedure (I/R or intra-cardiac injection), animals were given post-operative analgesics (buprenorphine, 0.1 mg/kg body weight) and allowed to recover until the experimental time points indicated where mice were then further analyzed, or tissue harvested. See final section for discussion of blinding and sample elimination considerations.

Infarct size and area-at-risk post-I/R was determined using triphenyl tetrazolium chloride (TTC) and Evans blue staining as previously described[75]. In experiments using immunosuppression via cyclosporine A (CsA), mice were anesthetized with 2% isoflurane inhalation to effect, and osmotic minipumps (Alzet, #1002) were implanted subcutaneously on the left lateral side of the mouse. Minipumps were loaded with CsA (Neoral, Novartis, NDC 0078-0274-22) dissolved in Cremophor EL (Sigma, #C5135) such that 15 mg CsA per kg body weight was delivered per day[76]. In experiments using $Kit^{MerCreMer/+}$ x *R-eGFP* genetic lineage tracing mice, tamoxifen was administered as previously described[42] via ad libitum feeding with pre-manufactured tamoxifen chow (tamoxifen citrate 40 mg/kg body weight per day, Envigo, #TD.1308603) for the time indicated in each individual experiment. For echocardiographic analysis of cardiac structure and function, mice were anesthetized with 2% isoflurane inhalation to effect and analyzed using a Vevo2100 instrument (VisualSonics) with an 18–38 MHz transducer as previously described[77]. See final section for discussion of blinding and sample elimination considerations.

**Histology and Immunohistochemistry**

Primary antibodies and dilutions used for immunohistochemistry are listed in Supplementary Table 1. For histological analysis, animals were anesthetized by isoflurane inhalation and sacrificed by cervical dislocation. The chest was opened, and the heart was flushed with cold cardioplegia solution (1 M KCl in 1x PBS) via cardiac apical insertion of a 25 gauge needle. The left atrium was cut to allow drainage of blood from the heart, and animals were briefly perfused with cold fixative (4% paraformaldehyde in sodium phosphate buffer, pH 7.4) through the apex of the heart. Tissues were excised, flushed with fixative, and incubated in cold fixative for 3.5 h at 4 °C with gentle rotation. Tissues were washed 3 times in cold 1x PBS and then cryopreserved by incubation in 30% sucrose in 1x PBS overnight at 4 °C with gentle rotation. Tissues were then embedded in TissueTek optimal cutting temperature (O.C.T) medium (VWR, # 25608-930) and flash-frozen at -80°C. Five-micrometer cryosections were cut using a Leica CM1860 cryostat.

Picrosirius red staining was performed with a kit from Abcam (ab150681) as per manufacturer's instructions. High magnification images of picrosirius red-stained hearts were captured at 200X magnification using an Olympus BX51 microscope equipped with a single chip color CCD camera (DP70) and DP controller software (Olympus America Inc., v3.1.1.). Border zone fibrosis was quantified as the percentage of picrosirius red-stained area over total tissue area analyzed as previously described[78].

All detection of genetic reporter-driven mTomato, RFP, GFP, or eGFP expression was performed using endogenous fluorescence without antibody labeling. Immunohistochemistry was performed on cardiac cryosections as previously described[42,74] with the following modifications (see Supplementary Table 1 for primary antibodies and dilutions. Alexa Fluor fluorochrome-conjugated secondary antibodies were used at a 1:200 dilution for visualization; Life Technologies). For immunohistochemistry using antibodies against PCM-1, antigen retrieval was first performed by incubation with 1% SDS for 5 min at room temperature with gentle rotation. Histological cardiac sections were washed thoroughly in 1x PBS before proceeding. To reduce non-specific background when using anti-mouse α-actinin on mouse tissue, cryosections were incubated with anti-mouse biotin (1:200) from the M.O.M. (Mouse-On-Mouse) immunodetection kit (Vector Labs, #BMK-2202) for 10 min at room temperature, followed by fluorophore-conjugated streptavidin. For immunohistochemistry using the collagen hybridizing peptide (CHP; 3Helix, #BIO300), a stock solution of 15 μm biotin-conjugated CHP was prepared according to manufacturer's instructions. The solution was heated to 80 °C for 5 min to denature the peptide, as previously described[53], followed by rapid cooling on ice and incubation on tissue sections overnight at 4 °C. Sections were then washed and processed for secondary antibody staining with fluorophore-conjugated streptavidin antibody as described above. Confocal microscopy and image acquisition were performed using a Nikon Eclipse Ti inverted microscope equipped with a Nikon A1R confocal running NIS Elements AR 4.50.

**Passive Force Measurements**
Tissue strips from the infarct region of the left ventricle were dissected using a Zeiss Discovery V8 dissection microscope. Tissues were cut into 3 mm (length) x 2 mm (width) strips, and 3-4 strips were cut from each infarct region of the left ventricle. These strips contained scar and a small region of border zone on each end. Tissue strips were maintained in M199 media (Corning Cellgro, 10-060-CV) with no supplementation throughout the procedure of force measurements. Tissue strips were attached to aluminum t-clips (Kem-Mil, #1870) and mounted onto a permeabilized muscle fiber test apparatus (Aurora Scientific, Model: 802D-160-322) initially set to zero tension. Cardiac tissue length was then increased 5% over 50 ms, held for 450 ms, then stretched again from 5% to 50% in intervals of 5% with no period of relaxation, and force was monitored using DMC v600A software (Aurora Scientific). Change in force was calculated as the difference between max force generated after the 50 ms pull and the minimum force achieved after each time period. Minimum force was calculated when the rate of force decay was zero by solving for the derivative of the best fit trend line, which was a 2nd degree polynomial equation.

**RT-PCR from Isolated Infarct Tissues**
Tissue strips isolated from the infarct region as described above were homogenized with a Precellys 24 homogenizer (Bertin Instruments #03119.200.RD000) and RNA was isolated by using the RNeasy fibrous tissue kit according to the manufacturer's instructions (Qiagen #74704). One microgram of total RNA was reverse transcribed using random oligo-dT primers and a Verso cDNA synthesis kit (Thermo Fisher Scientific # AB1453A) according to manufacturer's instructions. Real-time PCR was performed using Sso Advanced SYBR Green (BioRad # 1725274) according to the following program: one cycle of 95°C for 10 min, one cycle of 95°C for 15 s, 40 cycles of 95°C for 15 s, 57°C for 10 s, and 62°C for 30s, and one cycle of 62°C for 30s. *Gapdh* expression was used for normalization. Primer sequences used are included in Supplemental Table 2.

**Flow Cytometry and Cell Sorting**
For analysis of surface markers on MNCs, cells were resuspended in 1x HBSS supplemented with 2% BGS and 2 mM EDTA and incubated with fluorophore or biotin-conjugated primary antibodies (see Supplementary Table 1) for 20 min at 4 °C with gentle rotation. Cells were then washed twice with 1x HBSS. For detection of biotinylated antibodies, cells were incubated with streptavidin-conjugated BV421 (BD Horizon #563259) for 15 min at 4 °C with gentle rotation and then washed twice with 1x HBSS. Samples were analyzed using a BD FACSCanto running BD FACSDiVa V.8.0 software (BD Biosciences) and using the following laser configuration: Blue (488 nm), Yellow-Green (561 nm) and Red (635 nm). Analysis and quantitation was performed using FlowJo V.10 (Tree Star, Inc.).

For flow cytometry analysis of whole-heart cardiac macrophage or neutrophil content, single-cell suspensions were first prepared using enzymatic dissociation and trituration as described above. Animals were anesthetized by 2% isoflurane inhalation to effect and sacrificed by cervical dislocation. Hearts were rapidly excised and briefly rinsed in cold cardioplegic solution (1M KCl in 1x PBS) prior to enzymatic dissociation. The pellet resulting from dissociation was resuspended in 1 mL of red blood cell lysis buffer (150 mM $NH_4Cl$, 10 mM $KHCO_3$, 0.1 mM $Na_2EDTA$) and incubated at room temperature for 5 min. Samples were then centrifuged at 400 g for 10 min at 4 °C and resuspended in 1x HBSS supplemented with 2% BGS and 2 mM EDTA. Cells were incubated with fluorophore-conjugated primary antibodies for 20 min at 4 °C with gentle rotation, washed twice with 1x HBSS and analyzed using a BD LSRFortessa running BD FACSDiVa V.8.0 software (BD Biosciences) and using the following laser configuration: UV (355 nm), Violet (405 nm), Blue (488 nm), Yellow-Green (561 nm) and Red (635 nm) to detect fluorophore-conjugated antibodies and/or endogenous RFP and GFP signal from *Ccr2*-RFP x *Cx3cr1*-GFP mice. Analysis and quantitation were performed using FlowJo V.10 (Tree Star, Inc.).

For isolation of cardiac CCR2$^+$ or CX3CR1$^+$ macrophages by fluorescence-activated cell sorting (FACS), hearts from *Ccr2*-RFP x *Cx3cr1*-GFP mice at 7 days post-I/R injury were isolated and dissociated to single-cell suspensions as described above for isolation of CPCs, except that the digestion solution was made in DMEM + 2% BGS and 1% penicillin-streptomycin instead of HBSS. Isolated cells were sorted by FACS using a Sony SH800S benchtop cell sorter in a BSL-2 biosafety cabinet. Endogenous RFP and GFP fluorescence were detected using 4 collinear lasers and CCR2$^+$ (RFP$^+$ GFP$^-$) or CX3CR1$^+$ (GFP$^+$ RFP$^{+ \text{ or} -}$) cells were sorted into 1.5 mL Eppendorf tubes containing DMEM + 10% BGS and 1% penicillin-streptomycin. Cells were then cultured on isolated cardiac fibroblasts as described below.

**Cardiac Fibroblast Isolation**
Hearts were excised from ten 8-week-old male and female *C57/Bl6J* mice and the ventricles and septum were isolated, rinsed in ice-cold PBS and minced into small pieces using sterile micro-scissors. Tissue fragments were digested in DMEM + 2% BGS and 1% penicillin-streptomycin containing type 2 collagenase (100 units/mL; LS004177, Worthington, USA) for 20 minutes at 37ºC under gentle agitation. The digested tissue was triturated repeatedly to promote tissue dissociation. Dense fragments were allowed to settle for 2 minutes and the supernatant, containing the cardiac fibroblasts, was collected and spun at 1000 g for 5 minutes. The cell pellet was resuspended in 10 mL of DMEM + 10% BGS and 1% penicillin-streptomycin and kept on ice. This process was repeated 3 times until all the tissue was adequately digested. To remove cardiomyocytes and cell debris, cell suspensions were spun at 300 g, followed by centrifugation of the supernatant at 1000 g. The final cell pellet containing the cardiac fibroblasts was resuspended in DMEM + 10% BGS and 1% penicillin-streptomycin and pre-incubated on 0.1% gelatin-coated plates for 2 hours to allow fibroblast adherence before replenishment of the cell culture medium.

**Cardiac Fibroblast and Macrophage Co-Culture**
Isolated cardiac fibroblasts were split into 24-well 0.1% gelatin-coated plates at a seeding density of 15000 cells/well and allowed to adhere overnight. Macrophage subtypes (CCR2$^+$ and CX3CR1$^+$ cells) isolated as described above were then seeded onto these cardiac fibroblasts at a density of 10000 macrophages per 15000 fibroblasts. Control fibroblasts received an equivalent amount of culture media containing no macrophages. Adherence was verified the following day by fluorescence microscopy for RFP or GFP. A total of 4-5 biological replicates was performed per group. Cells were isolated after 72 hours later for mRNA quantification.

**mRNA Isolation and qRT-PCR from Cultured Cardiac Fibroblasts**
Total RNA was purified from cultured cells with TRIzol reagent (Fisher Scientific # 15596018) according to manufacturer's instructions. Two hundred nanograms of RNA was reverse transcribed to cDNA using the Verso cDNA synthesis kit (Thermo Fisher Scientific # 277.97). Quantitative PCR was performed using SsoAdvanced Universal SYBR Green Supermix (BioRad # 1725274) and assayed in duplicate, according to manufacturer's instructions in a CFX96 PCR system. Primer sequences are included in Supplementary Table 2. All data were normalized to *Gapdh* (verified to not deviate in samples).

**Macrophage Culture on Fibrillar Collagen Patches**
Pre-sterilized resorbable collagen membranes (Ace Surgical Supply #509-3040) were cut into circles of a uniform thickness and a diameter of 6 mm and then placed into 96-well plates. Macrophage subtypes (CCR2$^+$ or CX3CR1$^+$ cells) isolated as described above and cultured in DMEM + 10% BGS and 1% penicillin-streptomycin were then seeded onto the collagen patches at a density of 10000 cells/patch. Control patches were incubated in culture media without macrophages. Five biological replicates were performed per group. After 5 days in culture, fibrillar collagen assembly was analyzed by second harmonic generation microscopy using a Nikon A1R multiphoton upright confocal

microscope equipped with a tunable Coherent Chameleon II TiSapphire IR laser set to 840 nm. Three images were randomly taken per patch and assessed by a blinded investigator to select the representative images for each group.

**Statistical Information and Experimental Rigor (blinding)**
All statistical tests used and graphical depictions of data (means and error bars, or box and whisker plots) are defined within the figure legends for the respective data panels. Exact *n* values for all experiments with statistical analysis are included in the figure legends or within the figure itself. For comparisons between 2 groups, unpaired Student's 2-tailed t-test was performed as noted within figure legends. For comparisons between more than two groups, a one-way analysis of variance (ANOVA) with Tukey's or Dunnett's post-hoc test was performed as noted within figure legends. $p<0.05$ was considered as statistically significant. All data were found to follow a normal distribution as determined by the Shapiro-Wilk normality test ($\alpha=0.05$). For experiments involving I/R surgery, the number of animals that received surgery was determined based on prior experimentation in the lab, which demonstrated a peri-operative surgical mortality rate of 20%. Only animals that did not survive a given surgical procedure or were found at the time of I/R surgery to have incomplete reperfusion (failure of slipknot suture release) were excluded from analysis, otherwise no exclusions occurred. Randomization of mice within a group to receive a given surgical procedure (I/R vs sham) or treatment (saline vs cells vs zymosan) was not needed because the mice were genetically identical and were littermates, although equal sex ratios were maintained. Echocardiographic analysis, quantitation of $eGFP^+$ endothelial cells or cardiomyocytes, quantitation of cardiomyocytes in cell cycle, measures of fibrosis, measures of tissue passive force, in vivo and in vitro gene expression analysis, and analysis of collagen organization in culture were conducted by investigators blinded to experimental treatment or procedure. Quantitation of macrophage content by immunohistochemistry was performed using automated fluorescence threshold analysis in NIS Elements 4.50.

**Data Availability**
All raw data generated or analyzed in this study will be made available from the corresponding author upon reasonable request.
